# Super-resolution light-sheet fluorescence microscopy by SOFI

**DOI:** 10.1101/2020.08.17.254797

**Authors:** Judith Mizrachi, Arun Narasimhan, Xiaoli Qi, Rhonda Drewes, Ramesh Palaniswamy, Zhuhao Wu, Pavel Osten

## Abstract

Here we describe a new method, named LS-SOFI, that combines <u>l</u>ight-<u>s</u>heet fluorescence microscopy and <u>s</u>uper-resolution <u>o</u>ptical <u>f</u>luctuation <u>i</u>maging to achieve fast nanoscale-resolution imaging over large fields of view in native 3D tissues. We demonstrate the use of LS-SOFI in super-resolution analysis of neuronal structures and synaptic proteins, including cortical axons, dendritic spines, pre- and postsynaptic cytoskeletal proteins and postsynaptic AMPA receptors, in thick mouse brain sections. We also introduce an algorithm to determine the number of active fluorophore emitters detected, allowing the localization of individual molecules in LS-SOFI images. We conclude that LS-SOFI is a versatile method for fast super-resolution imaging from any tissue of the body using both commercial and custom LSFM instruments.

---

Light-sheet fluorescence microscopy (LSFM; also known as selective plane illumination microscopy, SPIM)^1,2^ has become a widely used imaging modality, supported by tissue-optimized clearing protocols and a number of commercial and custom-designed LSFM instruments^3^. While LSFM is a versatile method that can rapidly image entire 3D organs and tissues, such as entire mouse brains, there remains a considerable trade-off between spatial resolution and the tissue volume that can be visualized. To help to address this limitation, we combine the advantages of LSFM with super-resolution optical fluctuation imaging (SOFI) that calculates super-resolved images using the temporal correlation of fluorophores’ “blinking” – on/off cycling through distinguishable states with different fluorescence intensities – in an image time series^4-6^. In the current study, this on/off property is provided by naturally blinking Alexa fluorophores^7,8^ (Alexa-fluor^®^ Plus; ThermoFisher) conjugated to secondary antibodies used for immunolabeling in thick brain sections.

To establish the conditions of SOFI-based enhancement of LSFM images, we acquired image stacks of free Alexa-488-conjugated antibodies diluted in an agar block that was optically cleared with a modified CUBIC (mCUBIC) solution^9^. The acquired image stacks (time series of 1,000 images acquired at 10 msec per frame; total acquisition time 10 sec) were first deconvolved with a Gaussian point spread function (PSF) matching the light-sheet illumination^10^ and then processed by a modified SOFI algorithm to calculate the spatial distribution of the Alexa-488 fluorescence over time with 2^nd^, 4^th^, 6^th^, and 8^th^ order cumulants across the entire field of view (FOV) of 820 x 410 μm (Supplementary Fig. 1 and 2), achieving a final super-resolution output of 50 x 50 nm lateral pixel size in the 8^th^ order super-resolution LS-SOFI images (Methods).

While the unprocessed LSFM images showed no signal above noise (Fig. 1a,d), the higher order LS-SOFI images showed clear “spots” of blinking signals, suggesting that the LS-SOFI analysis can sufficiently enhance signal to noise ratio (SNR) of the LSFM image series to detect small clusters of or perhaps even single Alexa-488 fluorophores (Fig. 1b-c, d-h). We quantified the signal enhancement by measuring the full width at half maximum (FWHM) of the intensity spectra across a single fluorescence spot taken from the 1^st^ to 8^th^ order cumulant analysis, which revealed a marked, >5-fold improvement from the 2^nd^ to 8^th^ order cumulant analysis (Fig. 1i-j). Finally, as a control, we also collected several 1,000 image stacks from an agar block prepared without the addition of the Alexa 488-conjugated antibodies. As expected, processing these image stacks by SOFI did not reveal any spots of focal blinking signal, confirming the specificity of the Alexa-488 signal in the above LS-SOFI results (Supplementary Fig. 3).

**Fig. 1.**
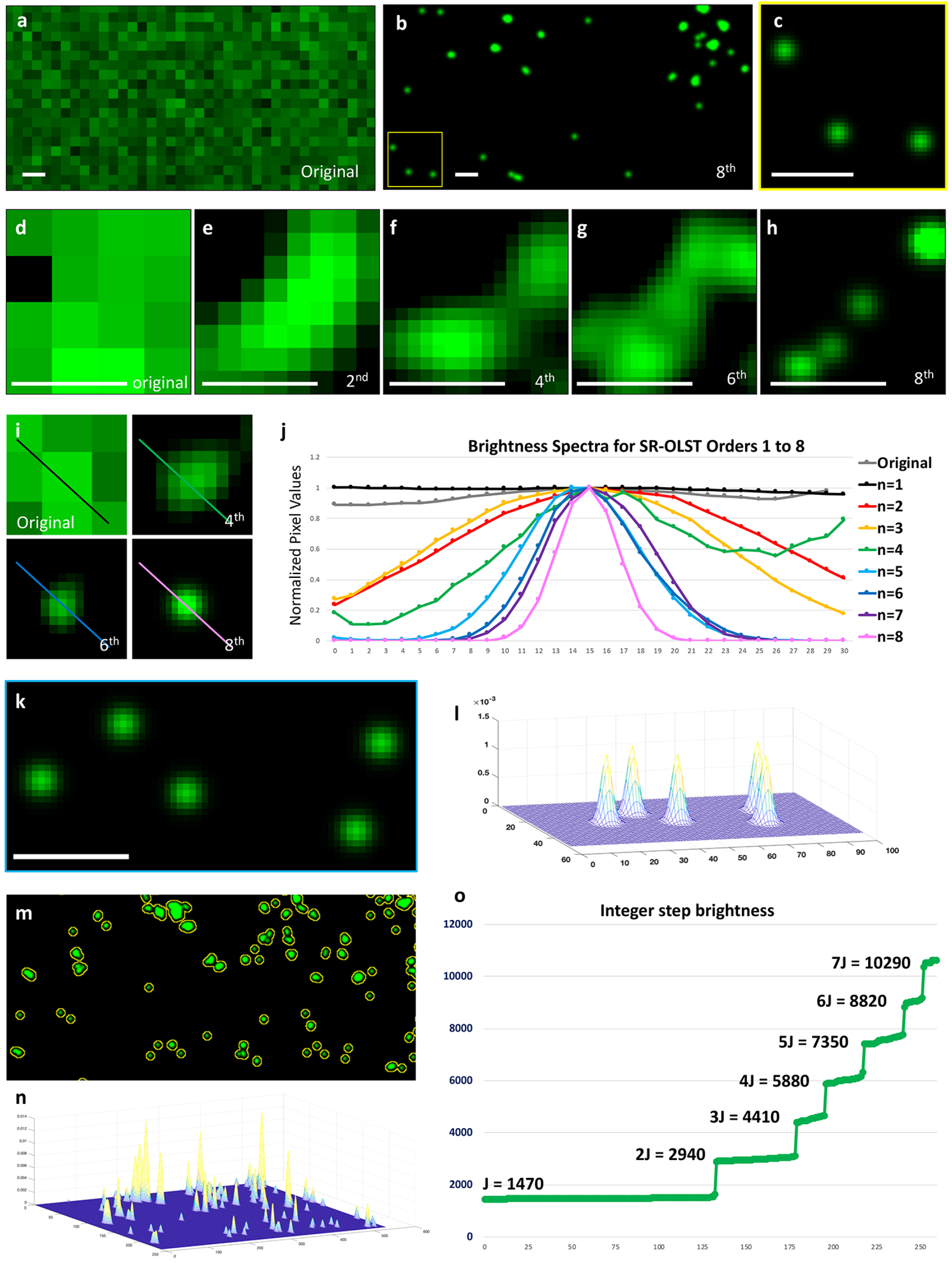
LS-SOFI-based detection of single and multiples Alexa-488 fluorophores. **a**, Single frame from a 10,000 time series taken from an agar block with diluted Alexa-488 fluorophores does not show any signal above noise. **b**, LS-SOFI 8^th^ order analysis of the same time series stack reveals distinct puncta of blinking fluorescent signals. **c**, Zoom-in view from (b) showing 3 fluorescence puncta of the same size, representing putative single Alexa-488 dyes. **d-h**, LS-SOFI 2^nd^ to 8^th^ order analysis reveals an incrementally more resolved image, from a lack of signal in the single frame (d), over 2^nd^ (e), 4^th^ (f), 6^th^ (g) and 8^th^ (h) order LS-SOFI revealing individual spots of fluorescence. **i-j**, Quantification of the signal to noise (SNR) improvement across a single fluorescence puncta (i) with LS-SOFI cumulant analysis of orders 1 to 8 (j). **k-l**, Another example of smallest fluorescence puncta (k) in the 8^th^ order analysis of uniform size, shape and brightness plotted in (l). **m-n**, Larger FOV shows fluorescence puncta in the 8th order analysis of various sizes, shapes and brightness plotted in (n). **o**, Plotting the total brightness of all spots of fluorescence detected generates discrete steps that are integer multiples of the first step derived from the smallest and uniform size spot. This indicates that LS-SOFI detects the number of active emitters as multiples of a single emitter that can be described as a constant “J”. Scale bar = 1 μm in all panels.

If the LS-SOFI analysis can detect single Alexa-488 fluorophores, we would expect the brightness of various sizes of individual signals (Fig. 1k-n) to be an integer multiple of the smallest signal representing a single fluorophore’s brightness. To test this, we next plotted the integrated total brightness over each fluorescence spot observed in the 8^th^ order LS-SOFI images from the agar-diluted Alexa-488 fluorophores. Strikingly, this analysis indeed revealed discrete steps of integer multiple values for all fluorescence spots (Fig. 1k-m), demonstrating that each spot’s brightness represents a multiple of the value of the lowest single emitter’s emission energy. Hence by approximating the value of a single emitter’s brightness, denoted here by the variable “J”, it is possible to detect single fluorophores as well as to estimate the number of active fluorophores in a given cluster region in the LS-SOFI data.

Having established the basic LS-SOFI imaging and computational parameters, we next applied this approach to visualization of neuronal structures – axons and dendritic spines – in mCUBIC-cleared 300 μm thick mouse brain sections from transgenic Thy1-GFP mice, in which GFP expression is targeted to and labels cellular morphologies of hippocampal and cortical pyramidal neurons^11^. The native GFP signal was converted to a “blinking” Alexa-based signal by immunostaining with anti-GFP primary antibodies followed by Alexa 488-conjugated secondary antibodies, before imaging selected areas of interest as 10,000 image stacks (10 msec frame rate, total acquisition time of 100 sec; Fig. 2 and Supplementary Fig. 4). In the first set of analyses, we compared LS-SOFI images produced by processing either 10,000 or 1,000 image stack taken from the same brain region, with the aim to empirically determine the effect of the expected 10-fold difference in the number of blinks that can be detected in these images. As shown in Fig. 2a-c, the structure of a single axon in the LS-SOFI images appears smooth in the SOFI-processed 10,000 image stack processed by 10^th^ order LS-SOFI, but is somewhat discontinuous (“patchy”) in the processed 1,000 image stack, demonstrating that in the shorter 1,000-frame series some regions appear to lack signal because fewer than 10 on/off cycles needed for the 10^th^ order LS-SOFI analysis were captured. Therefore, each preparation may need optimized conditions for detecting sufficient number of events to visualize the desired biological content.

**Fig. 2.**
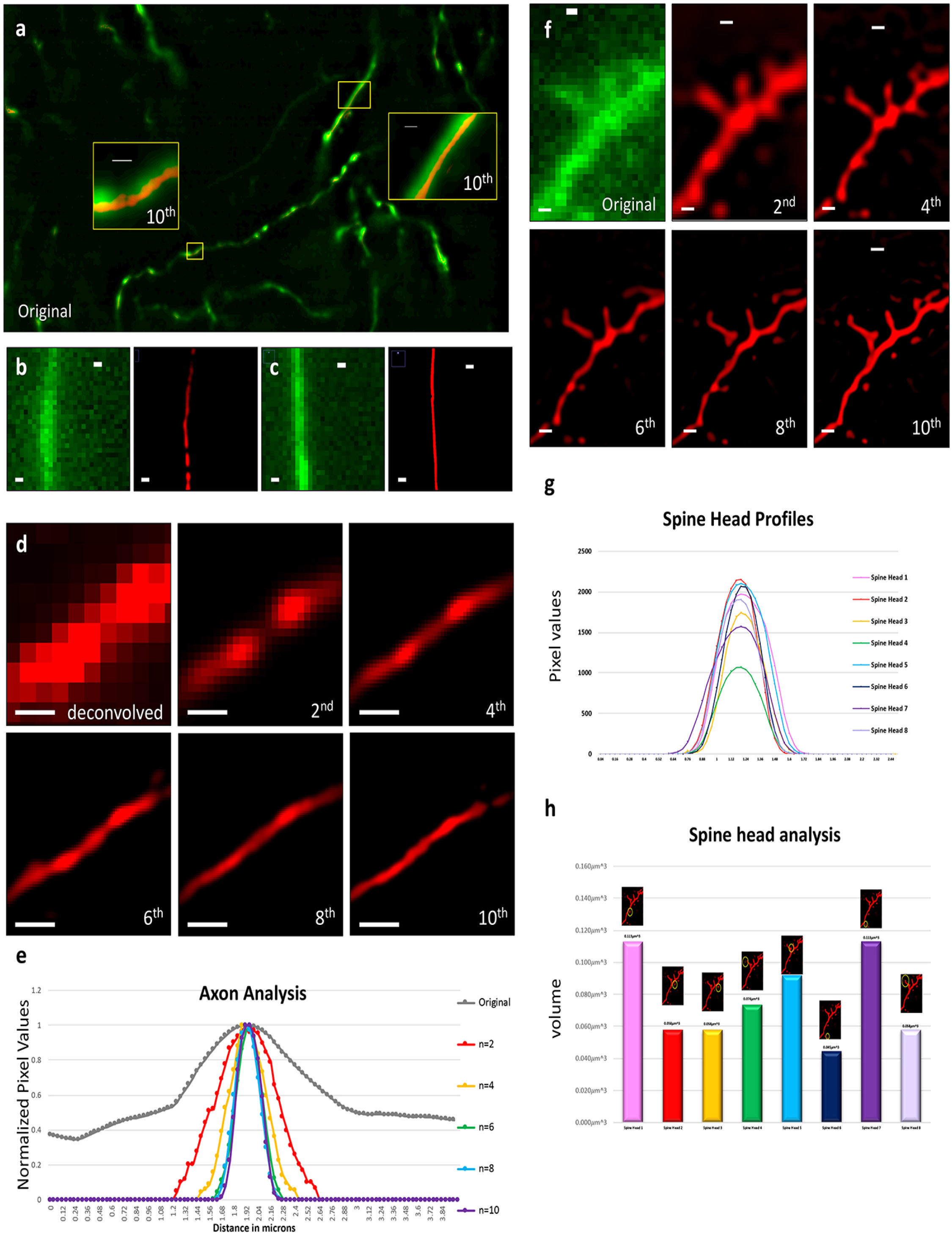
LS-SOFI enhanced imaging of neuronal morphology. **a**, Single image from a 10,000 time series taken from Thy1-GFP mouse cortex, with the two insets showing zoom-in views of axonal fibers labeled by GFP in green and overlaid narrower structures processed with 10^th^ order LS-SOFI of the full 10,000 image stack. **b-c**, Comparison of LS-SOFI 10^th^ order cumulant analysis of 1,000 (b) and 10,000 (c) frame time series stacks showing, respectively, an incomplete (patchy) and smooth continuous axon fiber. **d-e**, Quantification of the resolution of axon image with 2^nd^ to 10^th^ cumulant analysis (d) plotted as an analysis of cross-sectional resolution improvement (e). **f**, Progressive resolution improvement of cortical dendritic spines with 2^nd^ to 10^th^ cumulant analysis. **g-h**, Spine head morphologies derived in the 10^th^ cumulant analysis are analyzed by cross-sectional profiles (g) and spine head volumes (h). Scale bars = 1 μm in all panels.

Next, using the 10,000 frames image stacks, we compared the improvements in SNR for each step of higher order SOFI analysis of both axons and dendrites. First, GFP-labeled axon diameter comparison revealed a marked improvement in resolution, from >1 μm of LSFM resolution in single image frames to ~300 nm in both the 8^th^ and 10^th^ order SOFI with superresolution output of 50 x 50 and 40 x 40 nm lateral pixel size, respectively (Fig. 2d-e). Since the axon diameter did not further decrease in the 10^th^ compared to the 8^th^ order analysis, this indicates that the ~300 nm measurement represents the physical axon diameter in this image. This value also agrees well with axon diameter range of ~50 – 500 nm measured by electron microscopy (EM)^12,13^ and more recently using super-resolution STED (Stimulated Emission Depletion) imaging of GFP-labeled axons in mouse slice cultures^14^. Second, images of dendrites and dendritic spines showed a similar iterative resolution improvement with increasing order cumulant analysis, with the 10^th^ order LS-SOFI analysis of dendritic spines revealing a spine head cross-sectional diameter of 0.4-0.7 μm and an estimated average spine head volume of 0.076 μm^3^ (range 0.04-0.2 μm^3^) (Fig. 2f-h). These results are also consistent with EM measurements of spine size and spine head volume in the range of 0.06 to 0.07 μm^3^ in the mouse brain^15^.

In the next example of LS-SOFI analysis, we applied our technique to visualize pre- and postsynaptic sites immunolabeled with antibodies against the structural proteins Bassoon and Homer-1A, respectively. In these experiments, the brain sections were immersed in an oxygen scavenging buffer (glucose oxidase with catalase (GLOX)) shown previously to increase the on-off blinking of Alexa and other fluorescent dyes^8^ (Methods), and the data were collected as 1,000 image stacks (total acquisition time 10 sec at 10 msec per frame). Similar to what was seen in the axon and spine analyses, the 8^th^ and 10^th^ order showed comparable improvements in image resolution, revealing 2D profiles of both pre- and postsynaptic areas < 0.05 μm^2^ (Supplementary Fig. 5). To compare these data directly with an established super-resolution technique, we imaged the same tissue using SIM (Structured Illumination Microscopy), which revealed the same 2D profiles for both structures as measured by LS-SOFI (Supplementary Fig. 5), though notably the acquisition time of SIM was much longer, ~100 min compared to 10 sec of LS-SOFI. Finally, we note that the estimated sizes of the pre- and postsynaptic areas also agree with published measurements by super-resolution STORM using the same Bassoon and Homer1A immunolabeling^16^.

In the final set of experiments, we applied LS-SOFI to test whether the method can provide sufficient sensitivity to visualize the distribution of individual postsynaptic glutamate AMPA-type receptors labeled with antibodies against the GluA1 AMPA receptor subunit in thick brain sections prepared from the Thy1-GFP mice. Processing the 1,000 image stacks with 4^th^ order LS-SOFI analysis revealed a widespread distribution of fluorescence signals across the FOV, including along GFP-labeled dendrites and on neighboring tissue comprising unlabeled neurons and their processes (Fig. 3). In contrast, the 6^th^ order LS-SOFI analysis revealed focal blinking spots that in multiple cases aligned precisely along GFP-labeled dendrites, suggesting that these may indeed represent sites of GluA1 AMPA receptors. Furthermore, plotting the integrated total brightness over each fluorescence spot observed in the 6^th^ order images revealed again discrete steps of integer multiple values for the fluorescence spots, with each spot’s brightness representing a multiple of the value of the lowest single emission energy (Fig. 3c-d), suggesting that LS-SOFI can also localize individual postsynaptic AMPA receptors molecules in LSFM images of native brain tissue.

**Fig. 3.**
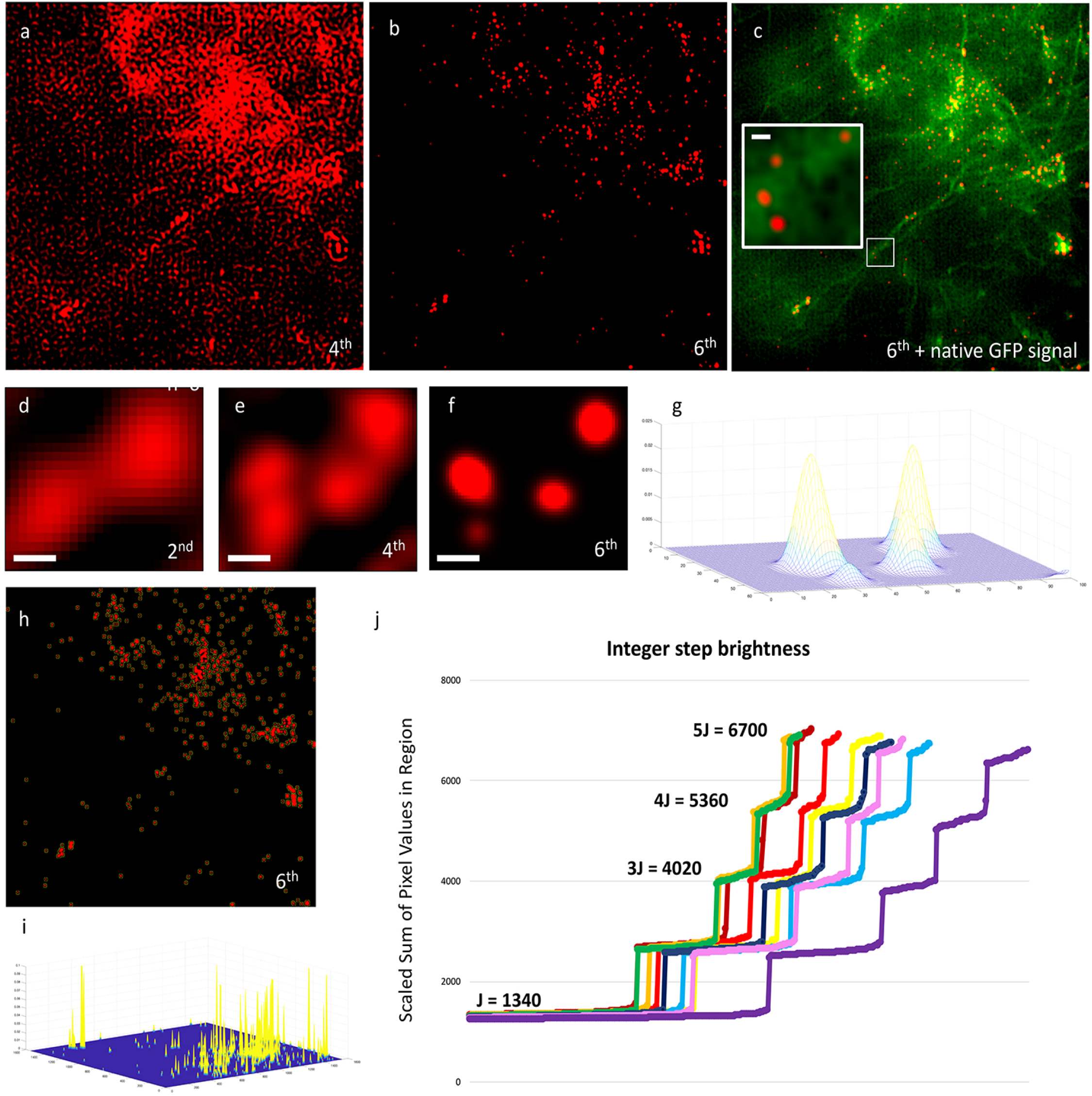
LS-SOFI enhanced imaging of AMPA receptor distribution. **a-c**, LS-SOFI of AMPA receptors (red) after 4th (a) and 6th order (b) cumulant analysis, with the 6^th^ order image overlaid over GFP-labeled neuronal morphologies (green). Zoom-in view in (c) shows several spots of fluorescence representing putative AMPA receptors. **d-g**, Progressive resolution improvement of spots of fluorescence with cumulant analysis of orders 2 to 6, plotted as analysis of size, shape and brightness in (g). **h-i**, Full FOV of fluorescence puncta in the 6th order analysis of fluorescence spots of various sizes, shapes and brightness plotted in (i). **j**, Plotting the total brightness of all spots of fluorescence detected in different images generates discrete steps that are integer multiples of the first step derived from the smallest and uniform size spot. This indicates that LS-SOFI detects the number of AMPA receptors as multiples of a single AMPA receptor labeled by Alexa-488-conjugated antibodies. Scale bar = 1 μm in all panels.

In summary, LS-SOFI is the first method to combine traditional LSFM and superresolution imaging, offering important advantages in terms of instrument versatility and low cost as well as speed of data acquisition and a broad range of tissue applicability compared to other super-resolution methods. First, LS-SOFI can be applied to time series images taken with any commercial or custom-built LSFM instrument in contrast to other super-resolution modalities, including STED, SIM and SMLM (Single Molecule Localization Microscopy), that all require specialized and expensive instruments^17,18^. Second, data acquisition times are much faster for LS-SOFI than for other super-resolution methods, as LS-SOFI images can be acquired in as little as 10 sec, while STED, SIM or SMLM methods require hours to generate super-resolution images from comparable FOVs. And third, LS-SOFI can be applied to any organ or tissue, such as thick brain sections in the current study, allowing versatile super-resolution analysis of structures and proteins in native 3D tissue environment in contrast to thin 2D sections used in other super-resolution methods.

## Acknowledgments

We thank

## Methods

### Tissue Preparation

Transgenic Thy1-GFP and wild type (WT) mouse brains were prepared as follows. Mice were deeply anesthetized with an injection of ketamine (100mg/kg) and xylazine (15mg/kg), transcardially perfused with chilled saline (0.9%NaCl) for 2 min, followed by fixative (4% Paraformaldehyde in 0.1M sodium phosphate buffer) for an additional 10 min. Following perfusion, the brains were carefully removed and placed in a tube with fixative solution for 48hrs.

For anti-GFP labelling in the Thy1-GFP transgenic brains, whole brains were delipidated using a modified adipo-clear protocol^19^ that will be described elsewhere (Zhuhao Wu; manuscript in preparation), after which the brains were sectioned into at 300 μm thick brain sections using a vibratome and stored in 0.05M phosphate buffer until immunolabelling using the iDISCO+ protocol^20^. Briefly, the tissue sections were incubated shaking for 30 min at room temperature (RT) in PTxwH buffer consisting of phosphate buffered saline, 0.1% Triton X-100, 0.05% Tween-20, 0.002 mg/ml heparin and 0.02% NaN_3_, followed by anti-GFP primary antibody incubation in the PTxwH buffer over-night (O/N) at RT, followed by 5 washed with PTxwH buffer for 1 min, 15 min, 30 min, 1 hr and 2 hrs, followed by secondary antibody in the PTxwH O/N at RT, followed by washes with PTxwH buffer for 1 min, 15 min, 20 min, 1 hr, 2 hrs and 4 hrs. Immunolabeling with anti-bassoon, anti-homer-1A and anti-GluA1 antibodies were done as with the anti-GFP antibody but without prior delipidation of the brain tissue. Before imaging, the brain sections were cleared with a modified mCUBIC solution for 24 hrs as described by us^9^.

### LSFM imaging

The 300 μm thick brain sections were carefully placed at the bottom of a 24-well plate, dried with a tip of a Kimwipe and embedded in 70 ^o^C 4% mCUBIC agarose and left to solidify. The agar block was carefully removed from the 24-well plate and the sample was glued to a glass slide and the slide placed in a 50-mm petri dish filled with mCUBIC buffer and placed in a custom built LSFM instrument called Oblique Light Sheet Tomography^9^. Regions of interest in the brain sections were imaged as 1000 or 10,000 frame time series stacks with FOV 820 x 410 μm.

### Data processing

The acquired image series were reconstructed as tiff stacks with a custom processing pipeline in Matlab which reads in the first frame of the raw data series and translates it into a MATLAB array variable, writes the array variable as tiff image type 16-bit unsigned integer, selects the next raw data frame in the input directory, and continues until image reconstruction of the entire dataset is complete^9^. Reconstruction of a 10,000 frame time series with a 410 x 820 μm FOV requires ~2 hrs.

Regions of high brightness, such as cell somas, within the selected field of view are manually segmented and excluded from the tiled image series for SOFI analysis. Either specific subregions or the entire FOV is tiled (200 x 200μm) across the entire image series for SOFI processing. After tiling, subpixel drift correction is initiated with a macro script in ImageJ plugin called AlignFullStack.ijm.ijm^21^, which performs robust and high speed sub-pixel alignment, assuming a reasonable reference point is specified and tissue drift is minimal. The aligned images of the full time-series are saved to a designated output folder for the aligned and tiled image series, together with a dataset file that contains the specifications of the alignment performed on the output image series.

The time series then undergo deconvolution for resolution improvement. The deconvolution software package is written into the general program that performs superresolution analysis on the tiled image series. The general call is with the bash script DeconSofiZframes.sh, which performs the necessary image analysis by calling the corresponding MATLAB program DeconSofiZframes.m. In both names the “Z” is replaced by the z-length of the time series stack. In the examples herein, Z has the value of 1000 or 10,000, reflecting the typical time series lengths of 1,000 and 10,000 frames respectively. The first stage of the DeconSofiZframes program begins by defining the tiled and aligned image series folder paths, and creating an output folder to which deconvolved and modified SOFI variables will be written.

The program reads the first tiled and aligned image series into an image stack and defines the PSF with which deconvolution with be performed. This PSF is selected from a set of customized model PSFs created with the Born & Wolf model^22^. The DeconSofiZframes.m program then calls a sub-function for sequential deconvolution called DeconToVar_100perRun.m. The program then writes the deconvolved stack to the output path designated by the X and Y sub-region of the original field of view that the tiled region spans. The deconvolved stack variable is then passed back to the DeconSofiZframes.m program to complete modified SOFI processing. The file path of the next tiled and aligned image series is then pointed to, and the process continues until the entire time series image set has been deconvolved. The final deconvolution program output is eight deconvolved, aligned, and tiled time series stacks of length Z, each with a 200×200μm field of view. When stitched together, this image series covers the entire 410×820μm original field of view with 10-pixel overlaps between tiles.

### Super-resolution by SOFI

The original version of the SOFI software package for super-resolution analysis has four main programs^6,23,24^. The first is “sofiCumulants” which performs pixel-wise analysis of cross-cumulants of the time-series image stack to generate a single array of these raw cross-cumulants. In its original edition, the “sofiCumulants” does not allow beyond 8^th^ order SOFI processing. The second is “sofiFlatten”, which flattens the cumulant value array found in “sofiCumulants” using the FWHM of a modelled PSF. This program also generates this modelled PSF. The new array of flattened cumulants produced by “sofiFlatten” replaces the original array created by the “sofiCumulants” program.

The third program “sofiParameters” uses the flat cumulants to estimate emitter parameters including on-time ratio, emitter density, and emitter brightness. The fourth program is sofiLinearize, which denoises and deconvolves the flat cumulant array, then linearizes the brightness by taking the n^th^ root of the array, since the brightness (ε) is raised to the n^th^ power in the n^th^ order cumulant function (Supplemental - SOFI proof). “sofiLinearize” then reconvolves the new array with a generated PSF. The final output is an array of linear cumulant values.

In our modified version of SOFI, the overall pipeline structure is largely maintained, however the programs themselves are changed in the modified SOFI software package to accommodate the specific requirements of LS-SOFI super-resolution analysis. The modified system begins when the deconvolution of the first tiled and aligned time series is complete. At this time, the deconvolved stack variable is passed back to the super-resolution analysis pipeline, and the modified SOFI package is initiated from within the DeconSofiZframes.m program.

The first program called from DeconSofiZframes.m in the modified SOFI package is sofiCumulantsHigherOrder_statusMonitor.m. This program is nearly identical to the original sofiCumulants program with two major changes that offer advantages with regard to LS-SOFI microscopy. The first is the ability to perform higher order SOFI analysis. The original SOFI software package only permits SOFI super-resolution up until cumulant order n=8. The current default settings for our pipeline perform super-resolution analysis for orders *n* = 1:10. The 10^th^ order modified SOFI images have a X,Y voxel size of 40×40nm.

To enable higher order cumulant analysis with our custom software program sofiCumulantsHigherOrder_statusMonitor.m, we created two additional sub-programs. The first is sofiGridsHigherOrder.m, which constructs the grid variable for higher order analysis. The n^th^ order grid variable is then passed back to the sofiCumulantsHigherOrder_statusMonitor.m program. We also modified the gpus.m software to create a corresponding software program gpusHigherOrder.m, which constructs higher order gpus and writes them to the designated higher order gpu folder for future use during modified SOFI analysis. The set of modified SOFI programs for higher order analysis are read from the folder directory SOFI_higherOrder, which is accessed directly from the processing cluster during super-resolution analysis. The second major modification to the sofiCumulants.m program is the user interface and status monitor output. The original sofiCumulants.m program had multiple calls to a graphic user interface output. In the case of LS-SOFI super-resolution processing, the software package is a completely automated system. The super-resolution processing pipeline is executed as a bash script that is submitted as a job to the institution’s cluster servers. In this application, therefore, the user interface and graphics initiated with the original software package are not practically suited to the LS-SOFI.

The modified SOFI processing software was also adjusted to designate the number of server nodes on which the program is run in parallel, as well as the memory usage of each node. This permits the super-resolution processing package to run in series, optimizing the process for faster overall analysis. If the default settings of the software package are selected, the programs are set to run within ten processing nodes, and each processing node is allocated 10GB of processing space. The settings can be altered as desired to suit the user’s preference, the total server space available, and desired processing time. After the n^th^ order cumulant values are calculated in the sofiCumulantsHigherOrder_statusMonitor.m program, they are passed to an n^th^ order array. This array and the higher order grid variable are then passed back to the DeconSofiZframes.m program and are saved to the designated output folder as MATLAB variables. Assuming the default settings are selected, each of the eight cumulant arrays calculated from each of the eight tiled and deconvolved time series are written to a corresponding output file, specifying the X,Y sub-region that the tile spans within the original field of view. For example, the raw cumulant array corresponding to the original pixel region 1:500, 1:500 is saved as “SOFIcumulantVar_x1to500_y1to500.mat”. The grid variable is written to the same folder with the name “SOFIgridVar_x1to500_y1to500.mat”. The cumulant analysis process repeats sequentially until all eight of the deconvolved sub-region tiles are complete. The raw cumulant and grid variables are passed on for final modified SOFI analysis.

After the raw cumulant analysis is complete and the cumulant and grid variables are written to the output folder, the raw cumulant array and the grid variable are passed on to final SOFI analysis from within the DeconSofiZframes.m program. This is executed in much the same fashion as the original SOFI software package, with modifications for higher order cumulant analysis and to predict and correct for the out of focus noise generated during LS-SOFI imaging.

The cumulant array and the grid variable are first passed to the sofiFlatten program, which calculates the flattened value array from the original raw cumulant data array. The new values are then written to the array designated by the “SOFI” variable, overwriting the raw cumulant data. The raw cumulants are flattened using the FWHM of a prediction PSF, which is generated from within the sofiFlatten program. Output variables are passed back to the DeconSofiZframes.m program and sofiParameters is called. sofiParameters takes the aforementioned array of flattened cumulants as its input and estimates the on-time ratio, the emitter density, and the emitter brightness. After the sofiParameters program is complete, the output variables are passed back to the DeconSofiZframes.m program once again, and the linearization step begins.

The linearization stage of the modified SOFI software package generates the final super-resolved images as an n^th^ order image array. This step is called from within the DeconSofiZframes.m program by executing the sofiLinearize program. This program denoises and deconvolves the flattened cumulant array. This program then linearizes the image brightness by taking the n^th^ root of the deconvolved cumulants. After the brightness linearization, sofiLinearize writes the new values to the “SOFI” array variable, overwriting the flattened cumulant values, and then reconvolves each of the *n* frames in the array.

Afterwards, the reconvolved frames are written to the “SOFI” variable, overwriting the previous values. This final “SOFI” variable array contains *n* images of modified SOFI orders 1 through *n* that constitute the final super-resolution images. The modified SOFI image array is once again passed back to the DeconSofiZframes.m program as the “SOFI” variable. A new output folder is created for the designated SOFI images of the sub-region tile field of view. The final image array is written to this folder as a MATLAB variable and as a series of *n* tiff images of type 16-bit unsigned integer. Upon completion of the super-resolution pipeline of all eight tiles, the program notifies the user with a message printed to the terminal that includes the output path location of the final super-resolution images. The DeconSofiZframes.m program then closes, returns to main bash script, and exits.

## Supplemental Fig. legends

**Supplementary Figure 1: LS-SOFI Software Schematic**

The LS-SOFI processing pipeline is composed of six steps, which can be called individually from their respective bash scripts or from a single automated call. These steps are: 1. Time series reconstruction, 2. Frame tiling, 3. Subpixel drift correction, 4. Deconvolution of aligned series, 5. Analyze Raw Cumulants, and 6. Complete Modified SOFI analysis. The final output of this software package is super resolution images of orders 1 through n.

**Supplementary Figure 2: Dependence of Number On/Off Cycles Captured on Camera Frame Rate**

The schematic illustrates the relationship between blinking rate, exposure time, and the number of on/off cycles detected in the acquired time series image stack. **a,** Exposure time equals the length of the fluorophore’s presumed dark state (10ms in this example). Each dark state spans more than 50% of at least one frame’s integration period, reducing the detected signal in that frame by at least 50%. Therefore, all five on/off cycles in this series are detected. **b,** Exposure time is much longer than emitter’s dark state. None of the dark states significantly reduce pixel value during the frame’s integration period, so none of the on/off cycles are detected. **c,** Exposure time is half the emitter’s dark state. As with (a), all five on/off cycles are recorded, but the image series is double the necessary length.

**Supplementary Figure 3: LS-SOFI of Agar Block**

**a-b**, Empty agar block imaged with LS-SOFI and processed with 8^th^ order cumulant analysis. 1^st^ order (a) and 8^th^ order (b) shown. No meaningful signal is apparent. (**c-f**) Full FOV of an agar block injected with Alexa 488 fluorophores imaged with LS-SOFI and processed with 8^th^ order cumulant analysis. 1^st^ order (**c,e**) and 8^th^ order (d,f) shown. **e,f** Zoomed regions outlined in (c,d) respectively. Individual emitters are visible in the 8^th^ order LS-SOFI image.

**Supplementary Figure 4: Additional axons**

**a-e,** Precise colocalization of super-resolution (red) and mesoscale (green) channels, acquireed from a 1,000 frame stack and distinguished by their blinking and non-blinking, respectively. **c-e,** Zoom-in of boxed regions in **(a). f,g,** Spectral analysis before (**f**) and after (**g**) super-resolution. The cortical axon’s diameter in this region is ~160nm, as shown. Samples were 300 μm thick coronal sections of a thy1gfp transgenic mouse brain, immunolabelled against GFP with Alexa Fluor 488. The 1,000 frame time series was acquired in ~10 seconds. 10^th^ order super-resolution images have a 40×40nm lateral pixel size.

**Supplementary Figure 5. Homer1 and Bassoon analysis**

**a-e,** Progressive resolution improvement with orders 1-10 cumulant analysis of post-synaptic homer1. **f,g,k,l,** LS-SOFI of a 1,000 frame time series, 410×820um FOV, ~10 second acquisition time, of a 100μm mouse brain section labelled against post-synaptic homer1 (f,g) and pre-synaptic bassoon (k,l) before (f,k) and after (g,l) 10^th^ order cumulant analysis. **h,m,** Areas of post- (h) and pre- (m) synaptic terminals. Average areas were ~0.074um^2^ and ~0.072um^2^ for post- and pre- synaptic terminals, respectively. The majority were smaller than 0.05um^2^. The region was imaged with confocal (i,n) and structured illumination (j,o) for validation.

