## Supplemental Figures for "Super-resolution light-sheet fluorescence microscopy by SOFI"

#### Supplementary Figure 1 : LS-SOFI Software Schematic

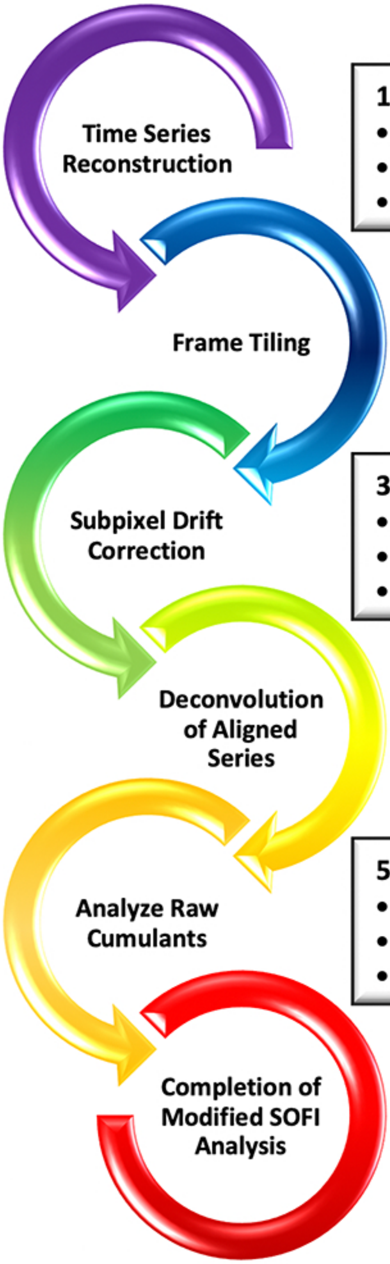


##### Super Resolution Processing Package

**1. Time Series Reconstruction**

- Call *datTotiff_singlestack.sh* => *datTotiff_singlestack.m*
- Reconstruct image time series from raw SR-OLST data
- Write as a 16-bit tiff image series

**2. Frame Tiling**

- Segment regions of high brightness from full field of view
- Call *cropStack_forAlignment.sh* => *cropStack_forAlignment.m*
- Divide entire FOV into 500x500xZ tiled image time series

**3. Subpixel Drift Correction**

- Call *AlignFullStack.ijm.ijm* to align the tiled image series
- Use stationary structures as reference points
- Write aligned series and alignment specs datasheet

**4. Deconvolution of Aligned Series**

- Call *DeconSofiXframes.sh* => *DeconSofiXframes.m* => *DeconToVar_100perRun.m*
- Selects PSF model => Deconvolves the tiled and aligned image series
- Writes the deconvolved image stack

**5. Analyze Raw Cumulants**

- Call *sofiCumulantsHigherOrder_statusMonitor.m*
- Default is $10^{th}$ order cumulant analysis, 40 x 40nm lateral pixel size
- Writes raw cumulant array and grid as .mat variable

**6. Complete Modified SOFI Analysis**

- Call *runJustSofi.sh* => *runJustSofi.m* for final modified SOFI steps
- Generate array of flattened cumulants, replace sofi variable
- Determine parameters => linearize cumulants
- Write super-resolution images

**Supplementary Figure 2**

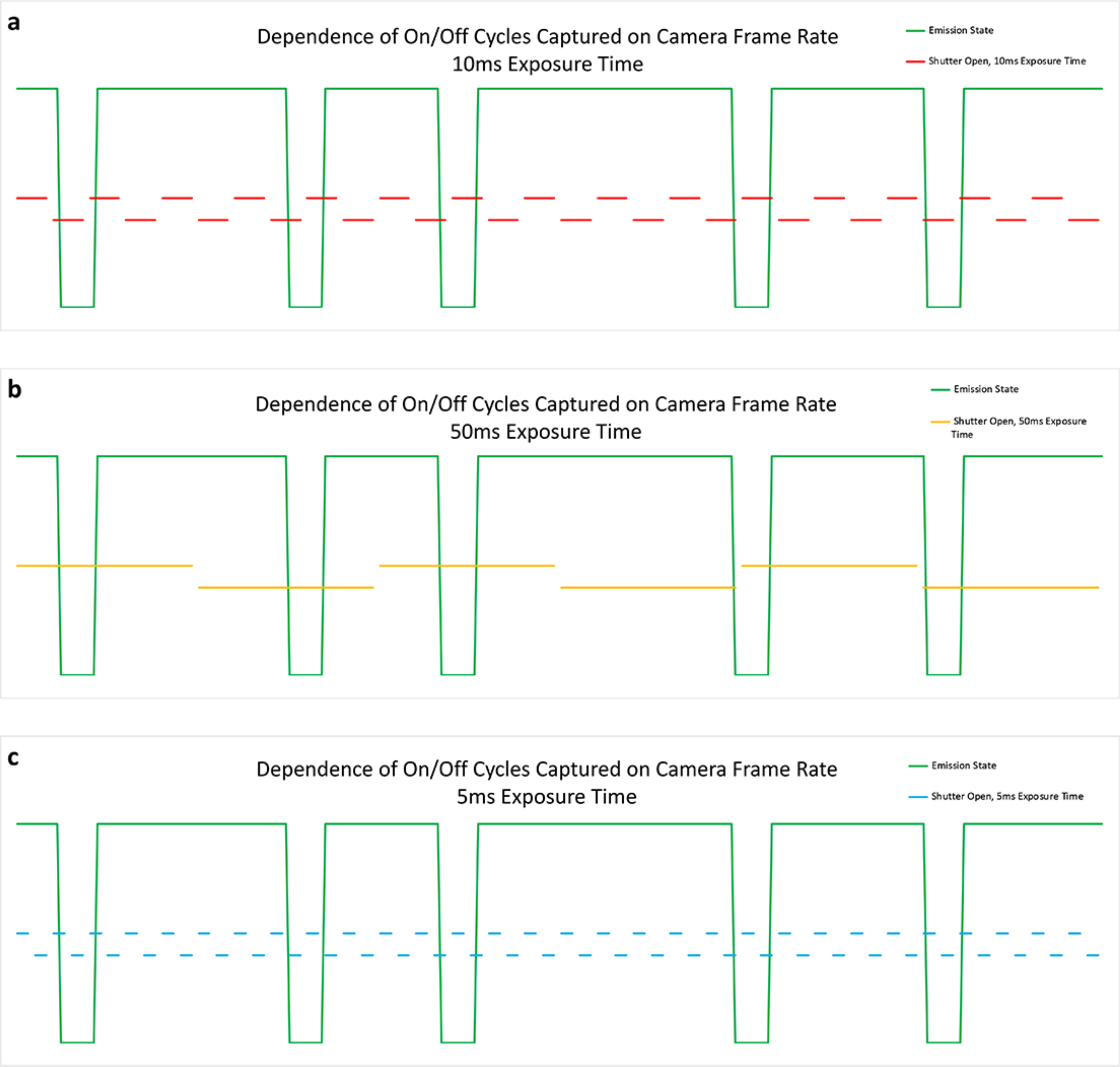

**Supplementary Figure 3**

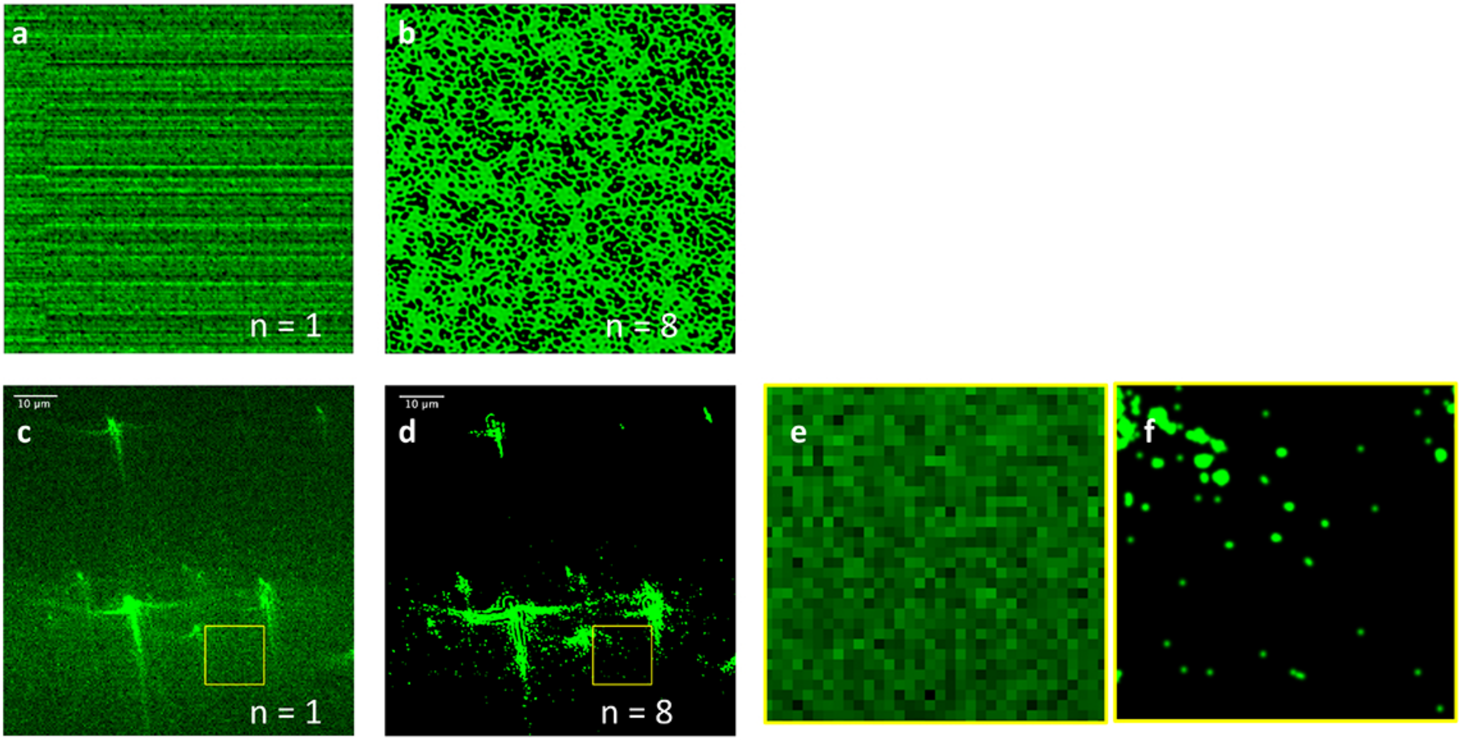

### Supplementary Figure 4

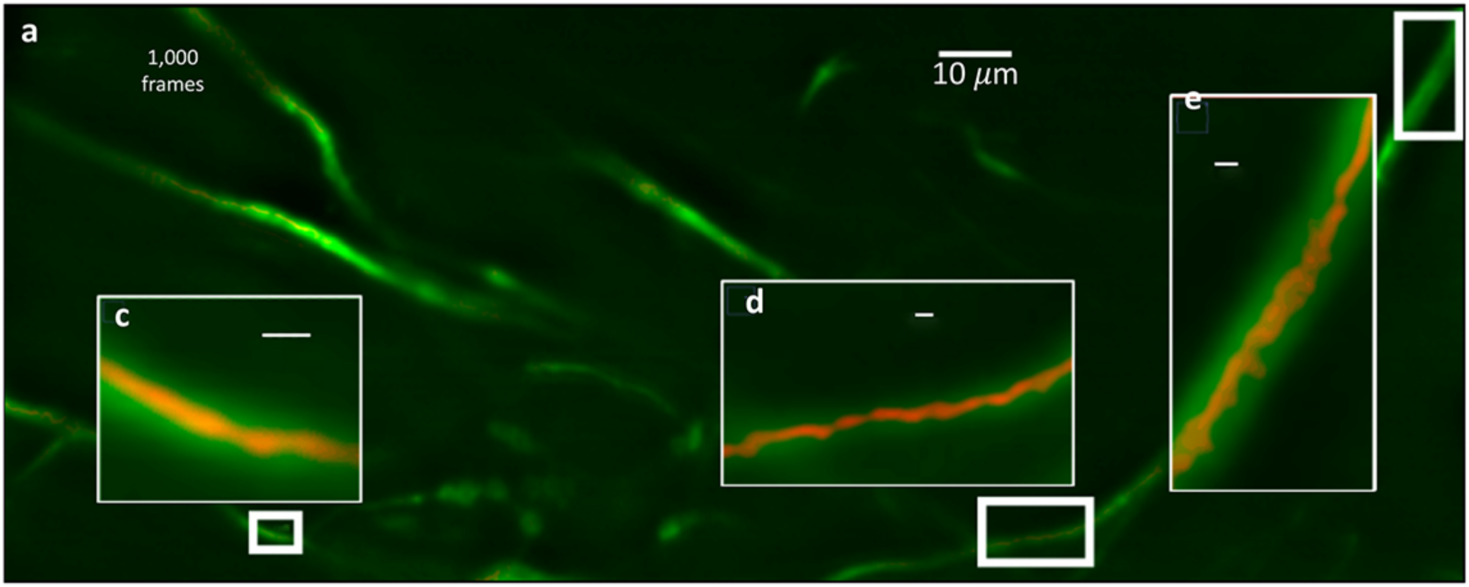


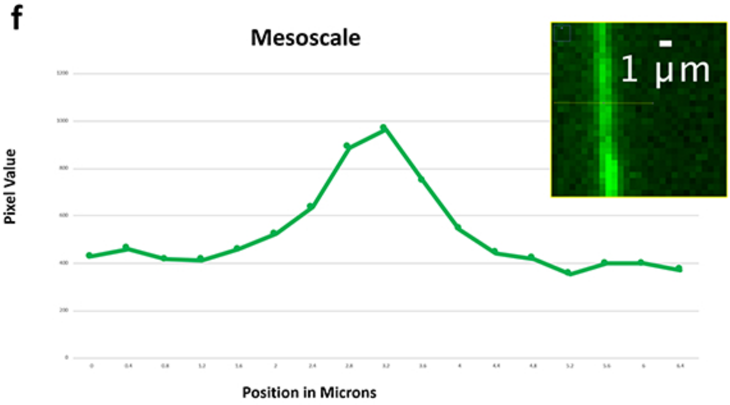


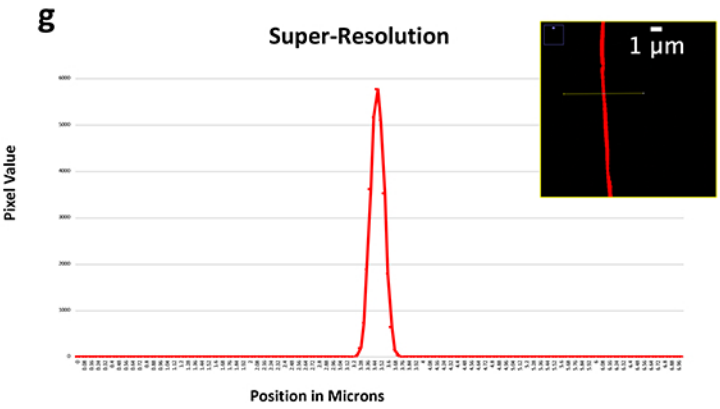

### Supplementary Figure 5

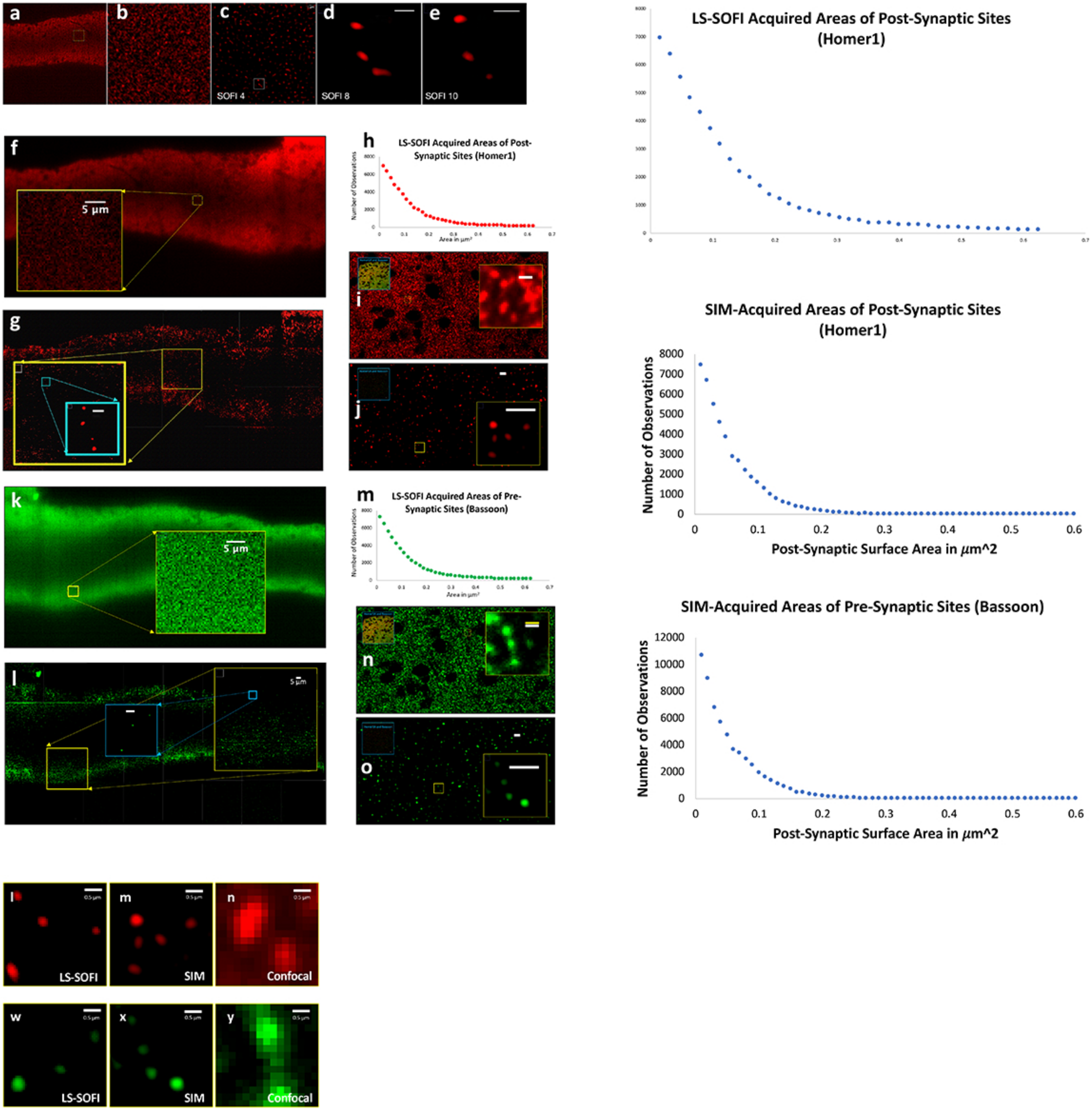
